# Cryo-EM structures reveal dynamic interplay of nascent chain-processing factors on the ribosome

**DOI:** 10.1101/312850

**Authors:** Sayan Bhakta, Shirin Akbar, Chiranjit Biswas, Jayati Sengupta

## Abstract

During protein biosynthesis in bacteria, one of the earliest phenomena that a nascent polypeptide chain experiences is the co-translational enzymatic processing. The event includes two enzymatic pathways, deformylation of the N-terminal methionine followed by methionine excision catalyzed by peptide deformylase (PDF) and methionine aminopeptidase (MetAP). The ribosome tunnel exit serves as the podium for recruiting proteins involved in maturation processes of the nascent chain. During the process, the emerging nascent protein likely remains shielded by the chaperone trigger factor (TF).

Here, we present the first cryo-EM structures of *E. coli* ribosome in complex with the nascent chain processing proteins. The structures reveal overlapping binding sites for PDF and MetAP when they bind individually at the tunnel exit site, where proteins L22 and L32 are identified as primary anchoring sites for both proteins. Interestingly however, MetAP has a remarkable ability of repositioning itself to adjacent locations in the presence of PDF and TF at the tunnel exit. Thus, our results disclose an unexpected scanning mechanism that MetAP adopts for context-specific ribosome association.

## Introduction

Ribosome not only serves as a platform for protein biosynthesis but also plays a crucial role in regulating various co-translational maturation steps related to protein biogenesis, including N-terminal enzymatic processing of the nascent chain (Kramer et al, 2009; Robinson et al, 2007). Emergence from the ribosome exit tunnel marks the onset of the maturation processes of the nascent polypeptide chain. The tunnel exit of the *E. coli* 50S subunit is surrounded by ribosomal proteins (L17, L22, L23, L24, L29, and L32) along with 23S rRNA helices (domains I and III) (Berk et al, 2006; Breiman et al, 2016). These proteins and rRNA domains around the tunnel exit serve as docking sites for co-translational enzymes and chaperones (Breiman et al, 2016).

In eubacteria, the nascent polypeptide chain encounters multiple members of a large repertoire of nascent chain processing enzymes, which primarily includes peptide deformylase (PDF), methionine aminopeptidase (MetAP/MAP), trigger factor (TF), signal recognition particle (SRP), SecYEG translocon, etc. How the enzymes modulate their anchoring positions on ribosome and accommodate one another in order to access the nascent chain properly is an interesting and yet unexplained question. Structural studies have demonstrated that SRP, TF and secYEG share overlapping binding sites on the ribosome (Halic et al, 2004; Halic et al, 2006; Schaffitzel et al, 2006), suggesting that the same platform is used for multiple functions.

Among all these factors, nascent polypeptide chain first confronts two proteolytic enzymes, PDF and MetAP (Meinnel et al, 1993). Protein synthesis in bacteria is initiated with N-formyl methionine (Giglione et al, 2015). As soon as the newly synthesized polypeptide chain reaches the extremity of the ribosomal exit tunnel, it first encounters PDF protein (essential in bacteria) that removes the N-terminal formyl group (deformylation). The PDF, a metallo enzyme, utilizes Fe (II) as the catalytic metal ion, which can be replaced with nickel or cobalt ion with no loss of activity (Ragusa et al, 1998; Rajagopalan et al, 1997). PDF subtype 1b interacts with the ribosome in *E. coli* (Bingel-Erlenmeyer et al, 2008).

Deformylation is a prerequisite for subsequent N-terminal methionine excision (NME) needed to produce diverse N termini in proteins for their appropriate function, targeting and eventual degradation. MetAP, existing in all kingdoms of life (Solbiati et al, 1999), is an enzyme that catalyzes the cleavage of first methionine from the growing polypeptide chain containing small and uncharged penultimate residue (Frottin et al, 2006). MetAPs also encompass metal binding site and can be activated by various divalent metal cations, but unlike PDFs, the physiological metal ions of MetAPs are still a matter of debate (Hu et al, 2007). *E. coli* possesses MetAP subtype1a comprising only the catalytic domain (Addlagatta et al, 2005; Lowther & Matthews, 2002; Lowther et al, 1999a; Lowther et al, 1999b).

Limited structural and functional studies related to the nascent chain interacting proteins in *E. coli* systems have been reported (Giglione et al, 2009; Giglione et al, 2015; Kramer et al, 2009). It has been established that the C-terminal helical extension of *E. coli* PDF is responsible for ribosome association. A crystal structure of the C-terminal helix of PDF in complex with the ribosomal large subunit has provided a model for PDF binding on the ribosome (Bingel-Erlenmeyer et al, 2008). This structure revealed that the ribosome-interacting C-terminal helix binds to a groove between ribosomal proteins L22 and L32, located next to the ribosomal exit tunnel, placing its active site accessible for interaction with the emerging nascent polypeptides. The ribosome interaction site of the MetAP has not been unambiguously identified thus far. However, several lines of evidence suggest that there is a MetAP association with the ribosome at the exit site of ribosome tunnel (Giglione et al, 2009). Biochemical and rigorous modelling studies have indicated the surface between L23 and L17 as MetAP interaction site where a positively charged loop of MetAP is the putative ribosome binding motif (Sandikci et al, 2013). It is evident that despite some progress, our understanding on how the tunnel-binding factors reciprocate with one another while simultaneously accessing the site remains obscure.

In this study, we present the first structural overview of the interactions of the nascent chain processing enzymes with the *E. coli* 70S ribosome (in the range of 10Å - 12Å resolution). By means of cryo-electron microscopy (cryo-EM) and 3D single-particle reconstruction techniques, three maps were generated where the nascent chain processing enzymes (PDF, MetAP) bind 70S ribosome individually (70S-PDF, and 70S-MetAP complexes) or together (70S-PDF-MetAP complex). The cryo-EM maps revealed overlapping binding sites of these proteins at the tunnel exit adjacent to the location of L22 and L32 proteins when they were allowed to associate with the ribosome independently (70S-PDF, and 70S-MetAP complexes). In contrast, when the proteins associate with the ribosome together (70S-PDF-MetAP complex), MetAP relocates at a neighbouring site close to L24, whereas PDF anchors at the same L22-L32 region. Another cryo-EM structure of a 70S ribosome in complex with PDF, TF and MetAP (70S-PDF-MetAP-TF complex), was also reconstructed where clear density corresponding to PDF and TF were seen, confirming that these two proteins can access ribosome simultaneously (Bornemann et al, 2014). An unambiguous localization of the MetAP density in this complex could not be achieved. However, substantial additional density was seen adjacent to the peptidyl-prolyl-cis/trans-isomerase (PPIase) domain of TF, which can be attributed to MetAP.

Taken together, our results reveal remarkable dynamic interplay of the protein biogenesis factors on the tunnel exit of the ribosome during post-processing of the nascent polypeptide chain, where ribosomal components also play an active role.

## Results

### PDF binding to the tunnel exit of the E. coli 70S ribosome

Purified *E. coli* 70S ribosomes were used to make the complex with *E. coli* PDF following co-sedimentation procedure (Fig. S1). A 3D cryo-EM map was generated for the PDF-bound 70S ribosome complex (map I, 70S-PDF complex, Fig1a). Additional density attributable to the PDF protein was clearly visible at the tunnel exit site in the 3D map (Fig.1a). Presence of PDF density at the tunnel exit was also visible in some of the views of the 2D class averages (Fig. S2a). Supervised classification was applied to remove the protein unbound population of 70S ribosome. In the resultant PDF-bound 70S ribosome map, the PDF density appeared to be very similar to the overall shape and size of the crystal structures (Becker et al, 1998) (Fig S3a). In agreement with previous reports, PDF is found to be accommodated in a groove created between large subunit proteins L22 and L32, adjacent to nascent polypeptide tunnel exit (Fig. 1d). Our 3D map manifested two prominent sites of PDF’s interactions with the 50S subunit. The protein’s C-terminal helix has been reported earlier as the binding partner of ribosome (Bingel-Erlenmeyer et al, 2008). In our 70S-PDF complex, the protein seemed to strengthen its hold on 50S subunit by employing another stretch of density protruding from its structure in addition to the C-terminal helix.

**Figure 1:**
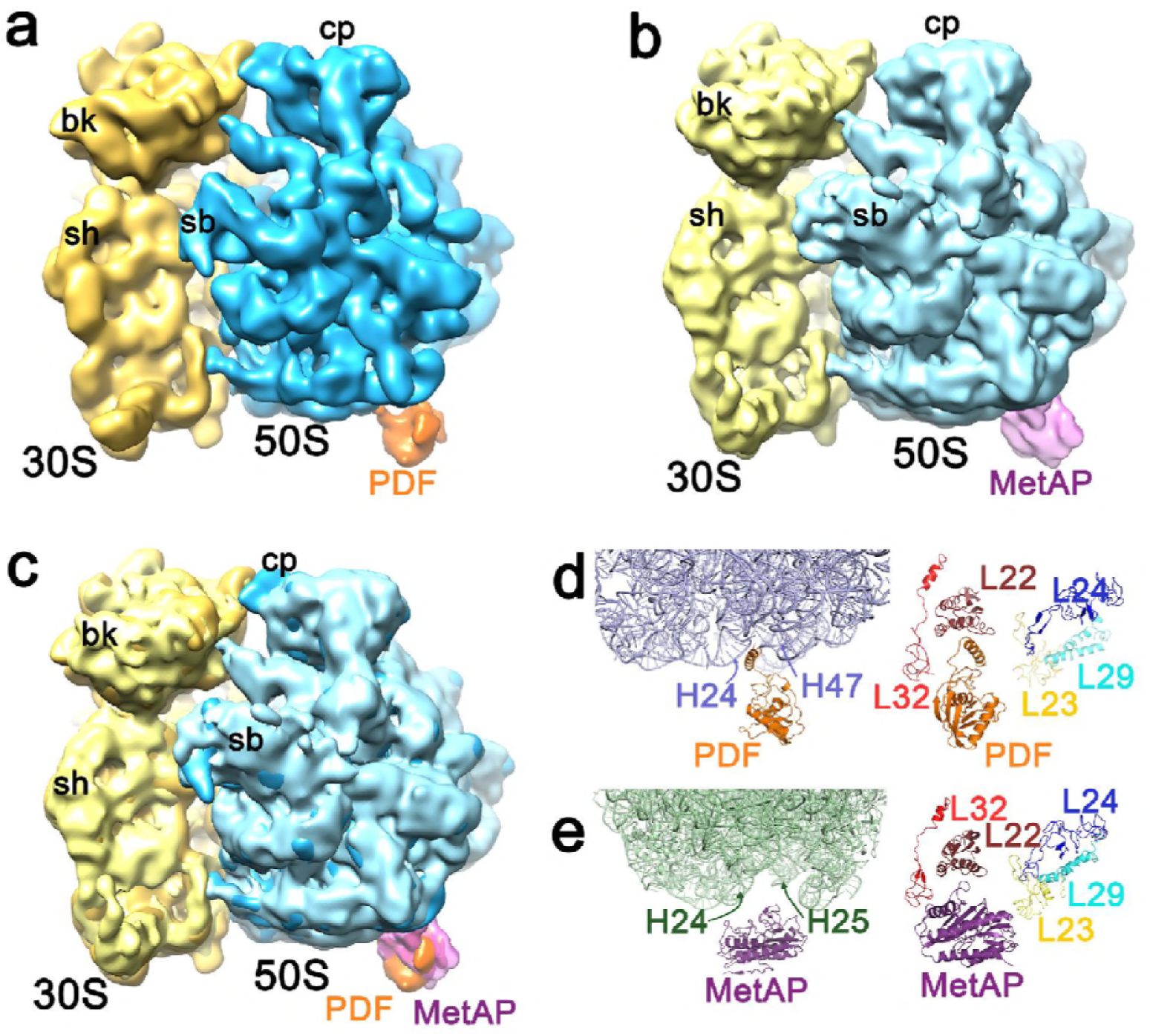
Visualization of PDF and MetAP at the ribosome tunnel exit when bound singly. a) Cryo EM reconstruction of *E coli* 70S ribosome-PDF complex showing the ligand at the tunnel exit site. The 30S subunit is shown in yellow, 50S subunit is shown in blue and the additional density for PDF is shown in orange. b) Cryo EM reconstruction of *E coli* 70s ribosome-MetAP complex. Density for MetAP is shown in magenta. c) Superimposition of 3D reconstructions of 70S-PDF and 70S-MetAP complexes showing common binding sites for PDF (orange, solid) and MetAP (Magenta, semitransparent). Landmarks for the 30S subunit: bk, beak; sh, shoulder. Landmarks for the 50S subunit: CP, central protuberance; sb, stalk base. d) Close-up view of quasi atomic model of 70S-PDF cryoEM map featuring regions of 50S subunit where PDF C-terminal helix (orange) and MetAP is located. C-terminal helix is seen lodged between the 23SrRNA (blue) helices H24 and H47 of domain I and III respectively which also positions the protein helix between ribosomal proteins L22 (chocolate) and L32 (red). e) Close-up view of quasi atomic model of 70S-MetAP shows location of MetAP on 70S is same as PDF. The interacting loops of MetAP encompassing several positive patches (Fig S3c) are seen stationed between 23SrRNA (green) helices H24 and H25 of domain I and seems to be in direct contact with L22 (chocolate).

It has been shown that empty 70S ribosomes are ‘unlocked’ and they spontaneously sample between ratcheted and non-ratcheted conformations (Valle et al, 2003). In agreement with the proposition, the dataset for control 70S ribosome (without any ligand) when subjected to 3D classification (Klaholz et al, 2004; Scheres, 2012) resulted into two distinct classes of ratcheted and non-ratcheted 70S ribosome maps (Fig. S3c-e). The topology of 70S ribosome in 70S-PDF complex when compared with the two control maps resembled non-ratcheted conformation (Fig. S3f,g), suggesting that PDF likely prefers to bind non-ratcheted conformation of the 70S ribosome. Interestingly, when the previously published atomic structures of ratcheted and non-ratcheted 70S ribosomes (Schuwirth et al, 2005) were overlaid, conformational changes were clearly detected, not only at the interface of ribosomal subunits, but also around the peptide tunnel exit (Fig.2a,b). A recent structure of the 70S ribosome in complex with secYEG with tunnel-bound peptide (PDB: 4V6M) revealed the interactions of L23 and L24 with the nascent polypeptide (Fig. 2c). The position of the nascent chain could be traced by aligning this cryo-EM model with other structures. Alignment of the tunnel-bound peptide with the ratcheted and non-ratcheted 70S ribosomes showed the relative orientations of the looped-out segments of proteins L23 and L24 with respect to the nascent chain (Fig. 2d,e).

**Figure 2:**
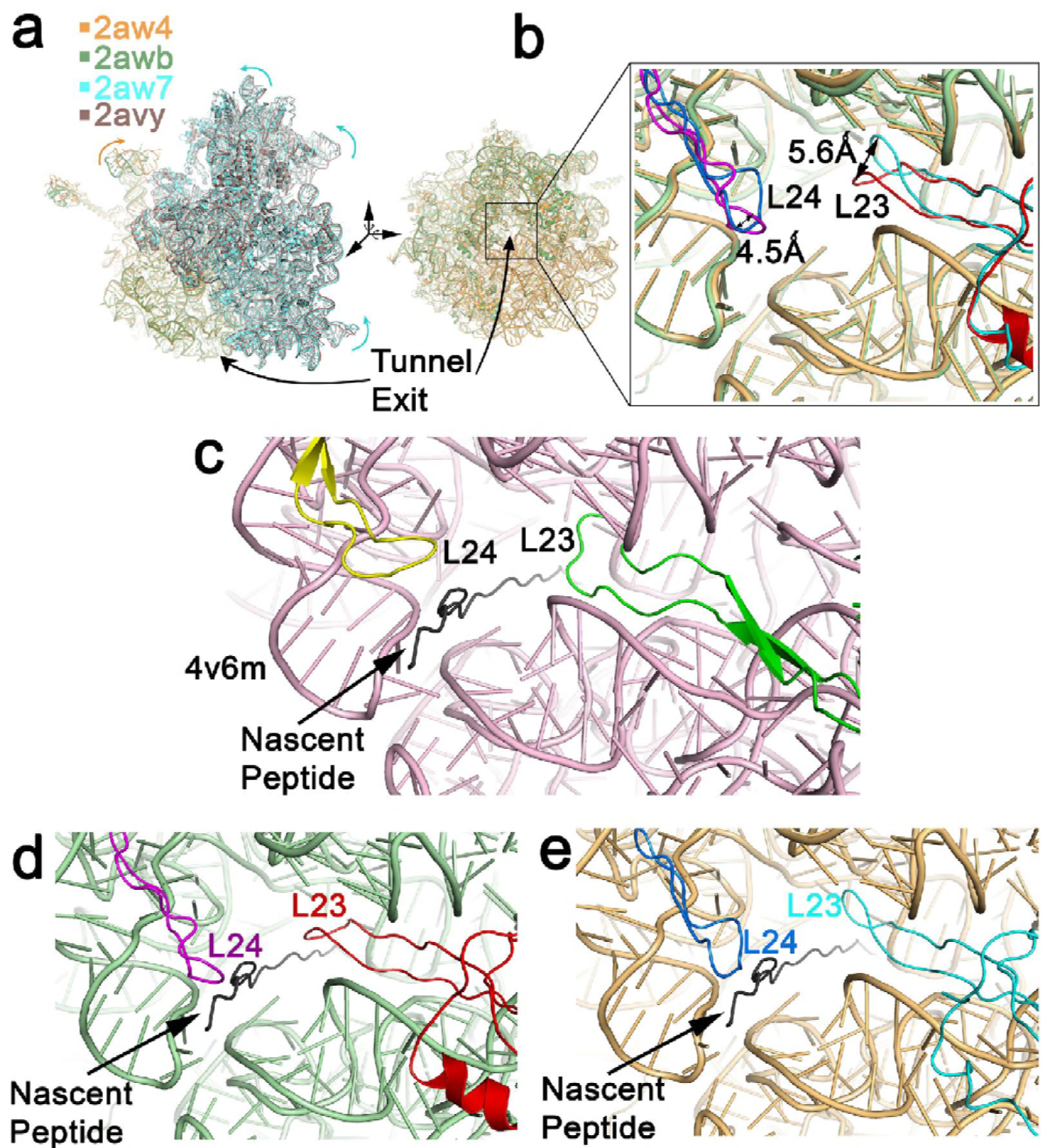
Effect of ratchet movement reflected at the tunnel exit. a) Superimposition of the ratcheted (PDB codes: 2avy, 2aw4) and non-ratcheted (PDB codes: 2awb, 2aw7) ribosome structures are shown. b) Close-up view of the tunnel exit shows significant conformational changes of the looped out segments of two ribosomal proteins (L23 and L24) around tunnel exit upon ratchet movement. c) Conformations of the L23 and L24 loops when nascent chain emerges from tunnel exit as revealed by the published structure. Alignment of the coordinates of 50S subunit containing nascent polypeptide chain (PDB code: 4v6m) traces interaction of nascent chain with L23 and L24 in case of (d) non-ratcheted (PDB code: 2awb) and (e) ratcheted (PDB code: 2aw4) ribosomes.

### The ribosome-PDF interactions at the tunnel exit

To gain further insights into PDF-ribosome interactions, quasi-atomic model of the 70S-PDF complex was generated using flexible fitting approach. Since the nascent chain processing proteins anchor on the peripheral surface of the large subunit, the cryo-EM map allowed molecular interpretation by using the atomic model reasonably well, despite the fact that the map was of moderate resolution.

In agreement with the previous structural study, the quasi-atomic model revealed that PDF anchors on the tunnel exit at the closest vicinity of L22 and L32 proteins (Fig. 1d). The C-terminal helix of PDF is the primary binding partner. The orientation of this ribosome interacting helix, however, is slightly different in the present 70S-PDF complex (Fig. S4a,b) as compared to that of the previously reported structure (Bingel-Erlenmeyer et al, 2008). The differences may be attributed to the fact that only the C-terminal helix was complexed with the 70S ribosome in the earlier structure, whereas the present structure represents the complex of the full protein with the 70S ribosome.

Closer look at the PDF-binding region identified the involvement of rRNA, as well as proteins around the tunnel exit in the interactions with PDF (Fig. 1d). Density corresponding to the full C-terminal helix (residues 147-168) is visible in our 70S-PDF complex map. While first half of the helix (residues 147-161) interacts with L22, the end part of the helix is accommodated in the groove made by the 23S rRNA helices H24 (domain I) and H47 (domain III). It should be noted here that 7 residues (aa 162-168) of the extreme end of the helix were not seen in the previous crystal structure (Bingel-Erlenmeyer et al, 2008), presumably due to the unstructured nature of that part (Fig. S4e,f).

### MetAP binds to an overlapping region of PDF binding site

Similar co-sedimentation approach was taken to make MetAP-bound *E. coli* 70S ribosome complex (map II, 70S-MetAP complex, Fig. S1). The 3D cryo-EM map (70S-MetAP, Fig. S2b) revealed extra density at the tunnel exit (Fig.1b) attributable to MetAP. The size of the density, representing MetAP (29 kDa), was elongated and flattened compared to the PDF density (19kDa) and resembled the MetAP crystal structures (Lowther et al, 1999a; Lowther et al, 1999b) well (Fig. S3b). Density corresponding to MetAP at the tunnel exit can be also detected in some of the views of the 2D class averages (Fig. S1b). Juxtaposition of PDF-and MetAP-bound maps (maps I and II) showed overlapping binding region for both proteins (Fig.1c), which is in contrast with the MetAP binding position proposed based on the previous docking study (Sandikci et al, 2013).

Although PDF and MetAP binding sites are overlapping, apparently the ribosomal binding partners of these two proteins on the tunnel exit are different. To determine the molecular details of interaction between the large subunit components and the MetAP protein, the quasi-atomic model of the complex was generated by flexible fitting approach. The atomic model allowed the interpretation of MetAP attachment site at molecular level.

Clear connections of MetAP-ribosome were seen at two positions (Fig.1b, Fig S3b), near the tunnel exit site on the 50S subunit, where the tunnel protein L22 resides. However, ribosome-MetAP connections were visible only at low contour levels, suggesting dynamic nature of the linker regions. Unlike PDF, where a dedicated C-terminal helix is the master regulator for ribosome association, MetAP possesses several flexible loop regions with positively charged residues likely responsible for ribosome binding (Fig. S4c,d). The involvement of Arg219 of one of the loops in ribosome association has already been demonstrated (Sandikci et al, 2013).

In contrast to the earlier report (Sandikci et al, 2013), MetAP positioned itself distantly from L17 and interacted with L22. With the positively charged loop regions, MetAP seemed to interact more profoundly with H24 of Domain III of 23SrRNA and looked closer to H25 (Fig. 1e). The ribosome interacting loop regions of MetAP seemed to come loose for better interaction. Difference in binding sites of the MetAP flexible loop regions within the immediate vicinity was predicted in an earlier study (Sandikci et al, 2013). Presumably due to the conformational dynamics of MetAP during ribosome binding, the overall MetAP density appeared to be incomplete. Moreover, when the ribosome conformation in MetAP-70S complex was compared with the ratcheted and non-ratcheted forms, it appeared that heterogeneous mixture of 70S ribosomes in both conformations was present. Thus, not only the dynamic nature of MetAP binding, but also heterogeneity in 70S ribosome conformations, seemed to contribute to lowering the resolution of the 70S-MetAP complex map.

### MetAP has multiple binding sites around the tunnel

Apparently, the nascent chain processing enzymes PDF and MetAP do not bind the tunnel exit very strongly when they bind individually. The observation that MetAP strongly competes with PDF, indicates that the primary binding site of MetAP overlaps with the binding region of PDF (Kramer et al, 2009; Sandikci et al, 2013). However, it has also been reported that the enzymes do bind simultaneously (Bingel-Erlenmeyer et al, 2008). To obtain insight into the interplay of the tunnel exit-bound proteins, we further reconstructed cryo-EM maps of two complexes: i) by incubating MetAP with the 70S-PDF ribosome pre-complex (map III, 70S-PDF-MetAP complex, Fig. 3, Fig. S2c) and ii) adding TF following 70S-PDF-MetAP complex formation (map IV, 70S-PDF-MetAP-TF complex, Fig. 4, Fig. S2d). Quasi-atomic models of the complexes were also generated.

**Figure 3:**
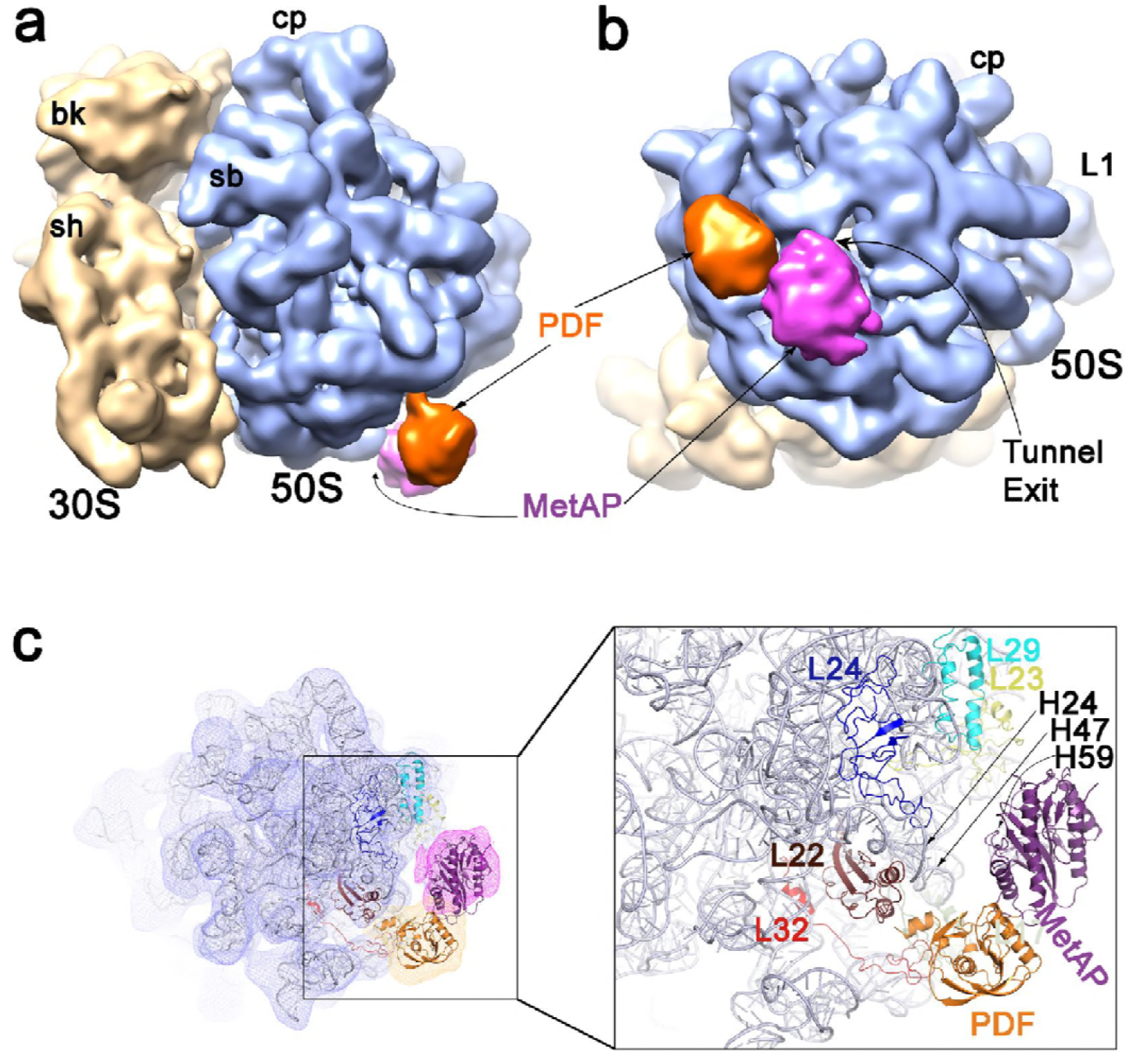
Cryo EM reconstruction of 70S ribosome-PDF-MetAP complex. a) Cryo EM reconstruction of *E.coli* 70S ribosome-PDF-MetAP complex where PDF and MetAP bind together at the tunnel exit. The 30S subunit is shown in yellow, 50S subunit is shown in blue and additional densities for PDF and MetAP are shown in orange and magenta respectively. MetAP relocates to an adjacent region in presence of PDF. b) Reoriented view from the nascent polypeptide tunnel exit site of 50S subunit show densities of MetAP (magenta) and PDF (orange) around the tunnel exit. c) A close-up view of the ribosomal architecture elucidating interaction zones between PDF, MetAP and 23SrRNA (steel blue), highlighting participant proteins L22 (chocolate), L32 (red), L23 (light olive), L29(cyan) and L24 (blue) surrounding the tunnel in different colours. Flexibility of the connecting loop of C-terminal helix allows rotation of the helix in opposite direction. Landmarks for 50S and 30S subunits are as in Figure 1.

**Figure 4:**
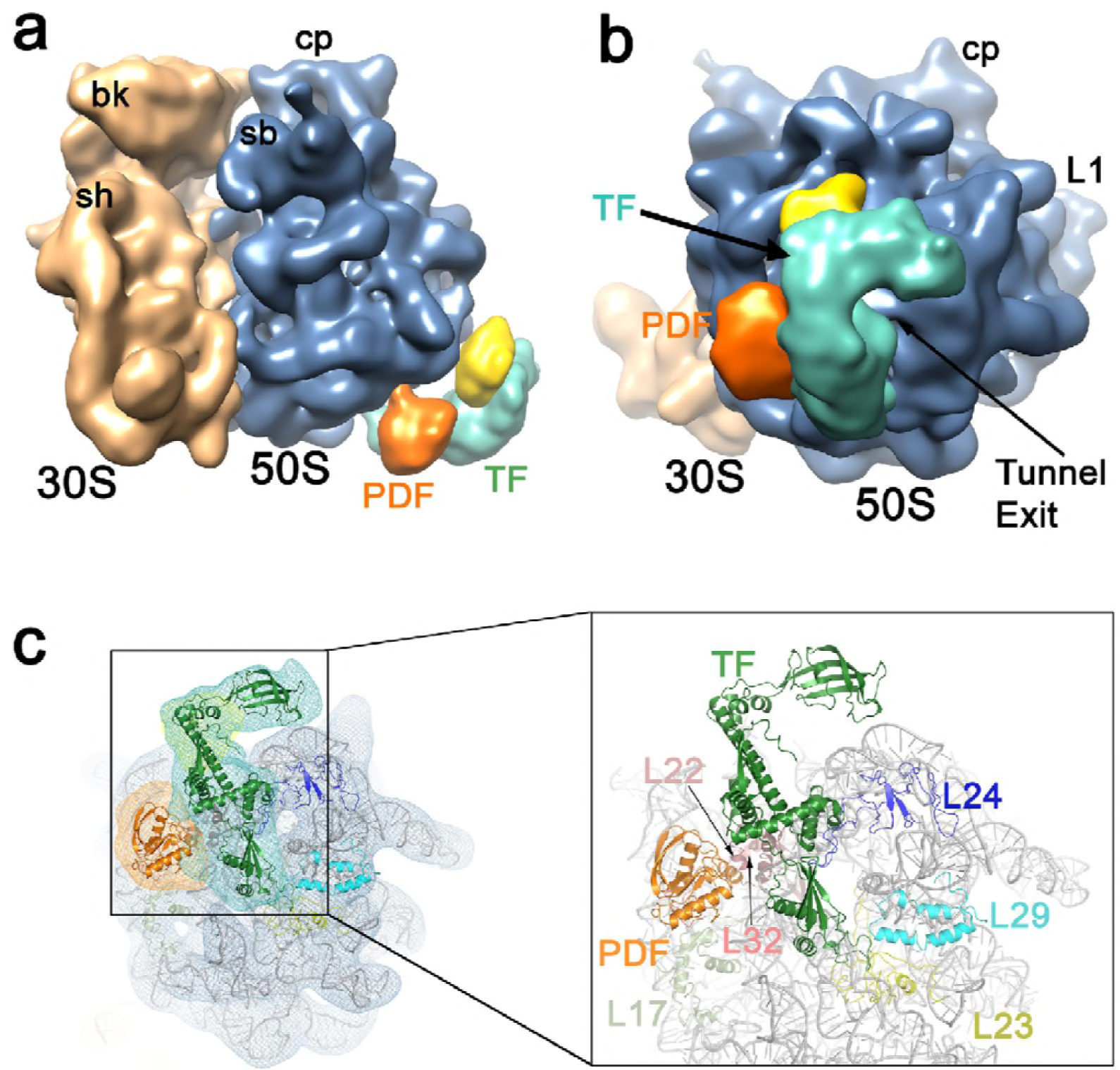
Cryo EM density map of the 70S ribosome-PDF-MetAP-TF complex. a) Cryo EM structure of *E. coli* 70S-PDF-MetAP-TF complex. 30S subunit is shown in wheat color, 50S subunit in blue shade, PDF in orange, TF in teal, and probable density of MetAP in yellow colours. b) Tunnel view of the structure showing densities corresponding to PDF and TF full proteins near the tunnel exit. The probable density of MetAP (yellow) is seen attached to the PPIase domain of TF and no connection with the ribosome is visible. c) A close up view showing interactions of PDF and TF with the ribosomal counter parts. PDF binds at the same close neighbourhood of the proteins L22 and L32. A loop of the ribosome binding domain of TF is interacting with L23.

It was clearly seen that while PDF occupied the same position as observed in the 70S-PDF complex, MetAP relocated at a position closer to L24 when compared to its position on the 70S-MetAP complex (Fig.3). Visualization of stronger density of the ligands in this complex (Fig. 3 a,b) suggested that the proteins mutually stabilized themselves on ribosome. Atomic model of the 70S-PDF-MetAP complex suggested that when MetAP binds adjacent to PDF, it almost entirely interacts with 23S rRNA. H59 of domain III seemed to be the predominant ribosomal partner of interaction. Other connections seemed to be with H50 and H47 of domain III of the 23S rRNA (Fig.3c).

In the 70S-PDF-MetAP-TF complex, PDF was found again in the same location and clear density corresponding to TF was also seen (Fig.4a,b). However, no obvious density corresponding to MetAP could be identified. In addition to the PDF density, strong density corresponding to the full TF was seen in the 70S-PDF-MetAP-TF complex map (Fig.4c). Complementarity of electrostatic surface potentials at the interface of PDF-MetAP (Fig.5a,b), as well as PDF-TF (Fig.5c,d) appeared to contribute in the stability of the binding of two proteins together.

**Figure 5:**
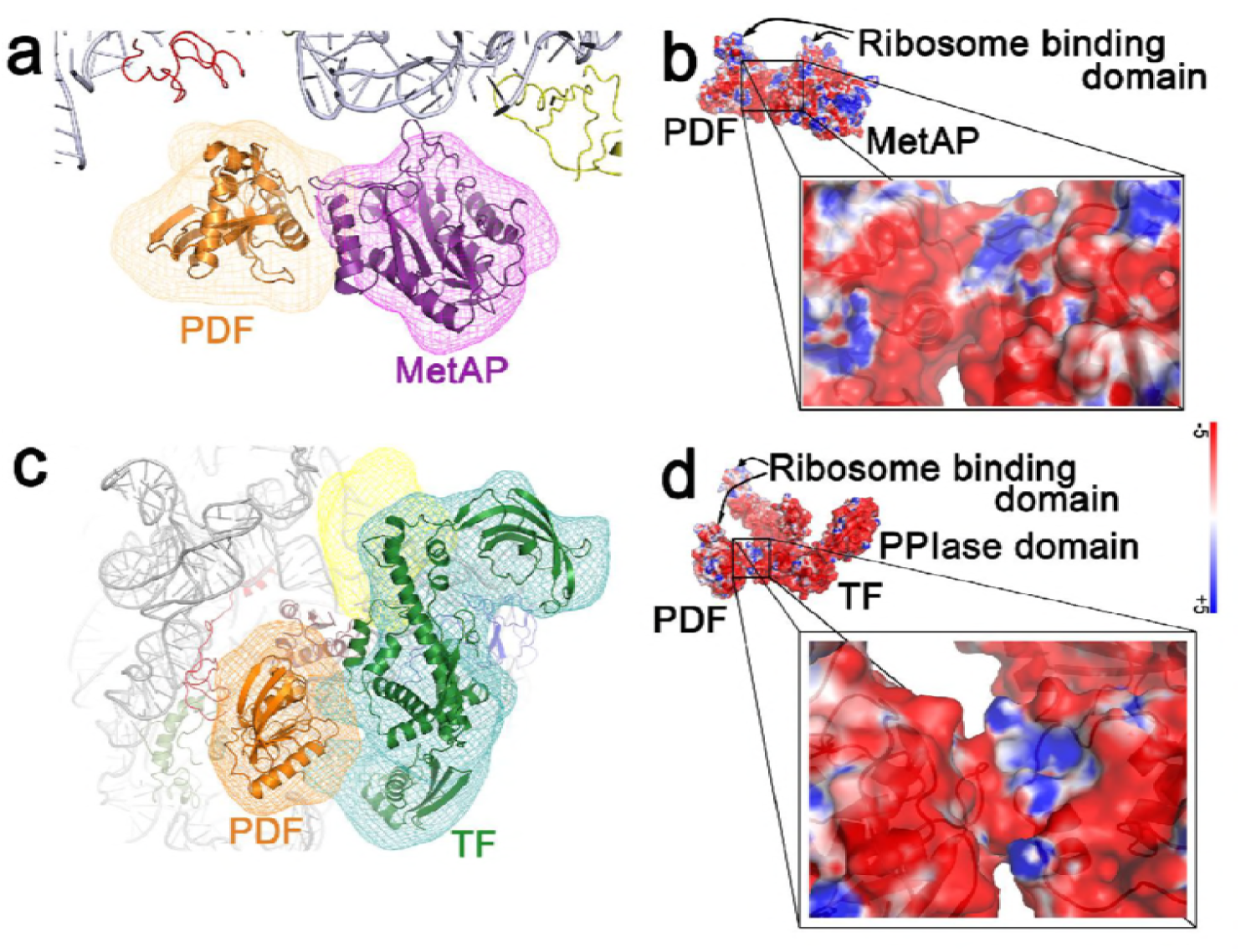
Interactions between cotranslational factors at the ribosome tunnel exit. a) Interaction between PDF and MetAP is shown with the fitted atomic models. b) Surface potentials (blue, positive; red, negative potentials), indicating complementary charge distribution at the interaction site of PDF and MetAP are shown. c) Interaction of PDF with TF is shown with fitted coordinates. d) Complementary surface potentials of PDF and TF at the connection between one arm of TF and a loop in the core structure of PDF.

Interestingly, a lump of additional density was found associated with the PPIase domain of TF (Fig.4a,b) in the 70S-PDF-MetAP-TF complex. Since there was no other ligand present except MetAP, we attributed this additional density to MetAP. This additional density was strongly associated with TF density, and although it was placed close to L24, no connection with ribosome could be detected. Thus, unambiguous identification of this density as MetAP was not possible.

We found that the position of the extended loop of L24 protein at the tunnel exit alters in the 70S-PDF-MetAP as well as the 70S-PDF-MetAP-TF complexes as compared to the 70S-PDF complex (Fig. S5a-e). It appeared that this loop of L24 in the altered position interacted with the emerging nascent chain (Frauenfeld et al, 2011). Notably, this region of L24 has been implicated to guide the nascent chain to PDF (Breiman et al, 2016).

### Rearrangements in PDF and TF conformations to accommodate other ligands

The position of PDF in the 70S-PDF-MetAP and 70S-PDF-MetAP-TF complexes was found to be similar to that of the 70S-PDF complex. However, some detectable differences in PDF conformations in the complexes suggest that PDF is also able to adjust its position to accommodate other ligands (Fig. 6). Although connecting density between PDF and ribosome tunnel exit was seen in both 70S complexes with additional ligands (70S-PDF-MetAP and 70S-PDF-MetAP-TF maps), density attributable to the C-terminal ribosome-anchoring helix (aa 145-168) was clearly not present at the same position (Fig.6g,k,h,l), as it was visible in the density map of the 70S-PDF complex (Fig. 6a-d). Interestingly, empty density was visible in the opposite side near the protein L22-32 cluster in both 70S complexes with additional ligands (Fig. 6e-i), where the helix (147-157) could nicely be accommodated (Fig.6f,j). The fitted position of the helix in the 70S-PDF-MetAP and 70S-PDF-MetAP-TF complexes is ~180° rotated compared to its orientation in the 70S-PDF complex. The feasibility of change in orientation of the helix is conceivable considering the long linker region connecting the helix with the rest of the molecule (Fig. S4a). Notably, although the crystal structure revealed helical structure of the C-terminal of isolated *E. coli* PDF, a previous study concluded a disordered nature of this part (Giglione et al, 2009; Meinnel et al, 1996). Moreover, the *S. aureus* PDF (PDF2) is perfectly superimposable with the *E. coli* PDF (PDF1) crystal structure except the C-terminal part, which is unstructured and ~90° rotated in *S. aureus* PDF.

**Figure 6:**
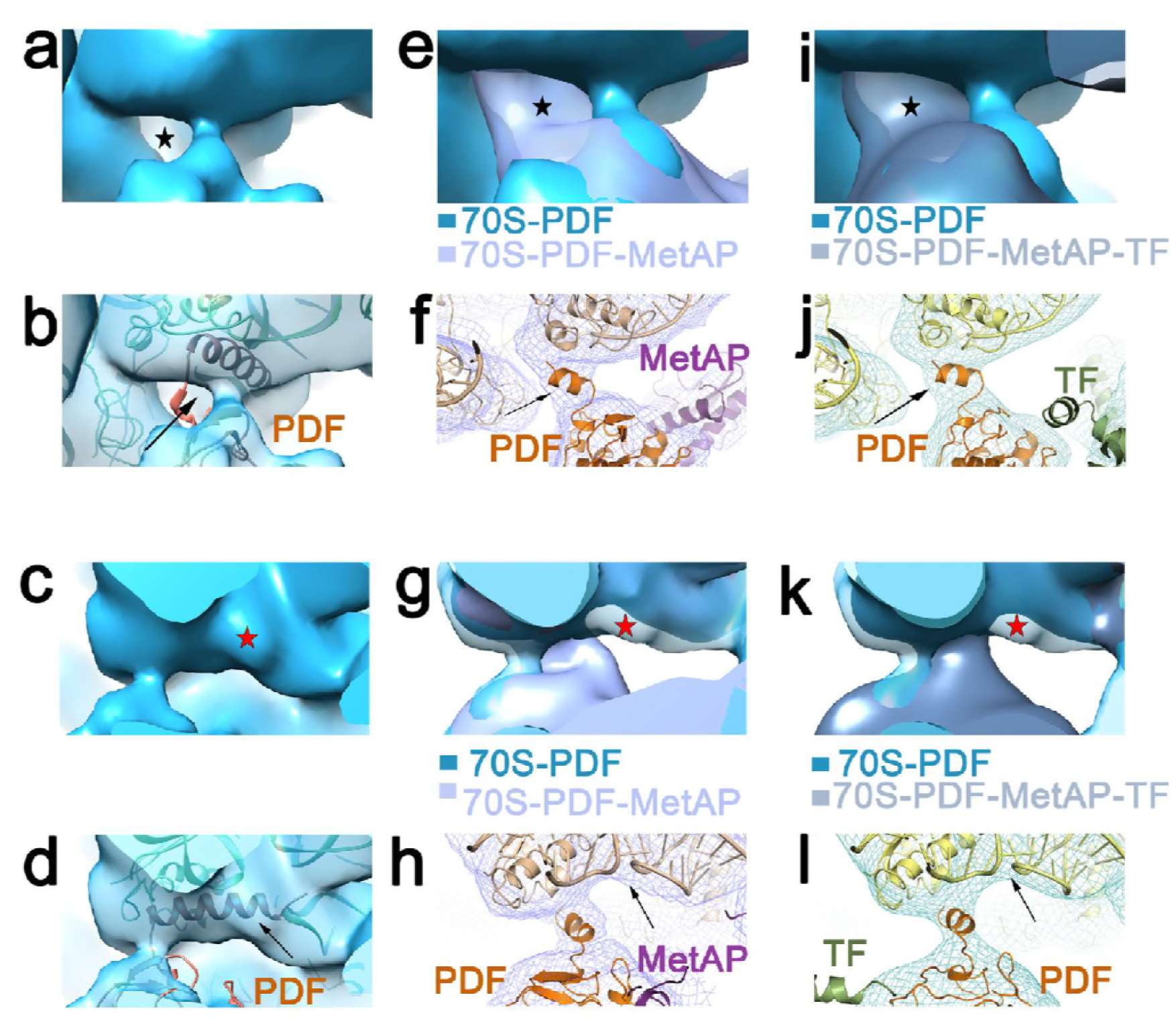
Reorientation of the C-terminal helix of PDF upon binding of MetAP or TF. Two close-up views of the PDF-ribosome interacting region, view 1 (a, b) and view 2 (c,d) of the 70S-PDF complex map show clear density (marked with red star in c) to accommodate full C-terminal helix (marked with arrow in d) is clearly available where the helix is placed in similar orientation as seen in previously published structure (Bingel-Erlenmeyer et al, 2008). Corresponding regions of 70S-PDF-MetAP (e,f) and 70S PDF-MetAP-TF (i,j) complexes are shown in view 1. Overlaying of the 70S-PDF complex map in (e,i) show the presence of additional density at the opposite side in both 70S-PDF-MetAP and 70S PDF-MetAP-TF maps (marked with black star) where majority of the C-terminal helix can be accommodated (f,j). View 2 shows the density corresponding to the helix in 70S-PDF complex is clearly missing in 70S-PDF-MetAP (g,h) and 70S-PDF-MetAP-TF (k,l). The observation hints to ~180° flipping of the C-terminal helix of PDF from its placement in 70S-PDF complex to its position in 70S-PDF-MetAP and 70S-PDF-MetAP-TF complexes.

It has been shown previously that in the presence of PDF, TF gets slightly shifted (Bornemann et al, 2014). In agreement with this observation, we found shifts and slight rearrangements in TF domains in the 70S-PDF-MetAP-TF complex (Fig. S5f,g) in order to avoid steric clash with the PDF, although the available coordinates of ribosome-bound TF could be fully accommodated into the density attributable to TF (Fig.4c).

### Discussion

One of the major findings that our study offers is the structural elucidation of MetAP relocalization in the presence of other ligands. Although such relocalization of ligand has not been commonly reported, similar large scale movement of bacterial initiation factor3 (IF3) along translation initiation pathway has been identified recently (Hussain et al, 2016). It has been shown previously that MetAP strongly competes with PDF (Sandikci et al, 2013), suggesting that the primary binding site of MetAP overlaps with the binding site of PDF, as we have observed when the proteins bind singly at the tunnel exit. A mechanism of action has been proposed (Giglione et al, 2009), suggesting that PDF, MetAP and TF bind simultaneously to the tunnel exit where TF is responsible to guide the nascent chain to the enzymes for their respective jobs. However, not every nascent protein needs chaperone assistance for folding (Anfinsen, 1973). In such cases, PDF and MetAP may bind sequentially to perform N-terminal methionine deformylation followed by methionine excision. Although the nascent chain masked by formylation does not have any role in MetAP binding (Sandikci et al, 2013), MetAP may have affinity to the unmasked nascent protein and following the release of PDF, MetAP binds at the same location to access the deformylated nascent chain. Variability in MetAP conformation, indicated in our study when binding singly at the tunnel exit, suggests its conformational freedom to locally scan the area for the nascent chain.

It has already been established that presence of nascent polypeptide chain in the peptide tunnel is not a prerequisite for binding of the ribosome-associated, co-translational nascent chain, processing enzymes (Bingel-Erlenmeyer et al, 2008; Sandikci et al, 2013). It may, thus, be speculated that ribosome conformational changes have the dictating role in recruiting these factors. A major conformational variability in 70S ribosome results from a built-in instability in the architecture of ribosome that allows it to switch between canonical and ratcheted conformations spontaneously (Spirin, 2002). In fact, during translation, ribosome continuously samples between ratcheted and non-ratcheted states (Frank et al, 2007; Valle et al, 2003). Upon ratchet-like movement, conformational differences of looped-out segments of tunnel exit-bound ribosomal proteins (particularly L23 and L24) (Breiman et al, 2016; Schuwirth et al, 2005) indicate a possibility of their guiding role in recruiting tunnel-binding factors. It has been proposed that the tunnel exit site offers entropic freedom to the nascent peptide to explore conformational space (Kosolapov & Deutsch, 2009). However, a more recent study has shown that ribosome influences the dynamics of a significant population of nascent chains to get spatially biased (Knight et al, 2013). We propose that proteins L23 and L24 likely have the role of arresting the conformational freedom of the emerging nascent chain and channelizing it to the active site of PDF.

Binding of PDF marks the onset of protein biogenesis. When MetAP sits next to it, they seem to mutually stabilize binding of one another. In the 70S-PDF complex, the ribosome-interacting C-terminal helix is positioned in one direction which rotates when other ligands such as MetAP and TF approach. It may be speculated that in the 70S-PDF complex the C-terminal helix is placed in such a way that makes its active site accessible to the nascent chain emerging out of the tunnel. It is interesting to note that unlike MetAP binding, PDF on a ribosome is restricted to a particular location, which can be attributed to the fact that PDF is always the first enzyme that a nascent chain encounters while leaving the tunnel. When present together, PDF sits on a ribosome, carries out deformylation, and when MetAP approaches, PDF readjusts itself, which results in passing the deformylated nascent chain onto MetAP for further processing. The RNA affinity of MetAP allows it to bind both sequentially and simultaneously with PDF to access the tunnel exit. It is possible that L22, which reaches to the interior of the tunnel, recognizes the requirement and authorizes the binding of co-translational proteins to be of either sequential or simultaneous in nature and, consequently, the guiding of L23 and L24 for extending or retracting their loops.

A recent study showed that association of TF and PDF with ribosome can occur simultaneously with an altered arrangement in TF (Bornemann et al, 2014). This study also showed that binding of a trigger factor to ribosome is not inhibited by MetAP. Consistent with this result, our 70S-PDF-MetAP-TF structure provides a structural elucidation that, indeed, TF shifts from its usual position (seen in previous TF-bound ribosome structures) to accommodate itself when PDF is already present. Superimposing available TF crystal structures showed considerable flexibility of the arms and PPIase domain of TF. Based on our results, we propose that for nascent proteins that need intensive care to choose productive folding pathway, the nascent chain processing protein trio (PDF, MetAP and TF) provides an almost sealed cavity to assist proper folding (Fig. 7). Taken together, a consortium of all these co-translational factors and chaperones over the tunnel exit site serves a greater purpose and singles out a crucial role of protecting the vulnerable nascent chain at all cost.

**Figure 7:**
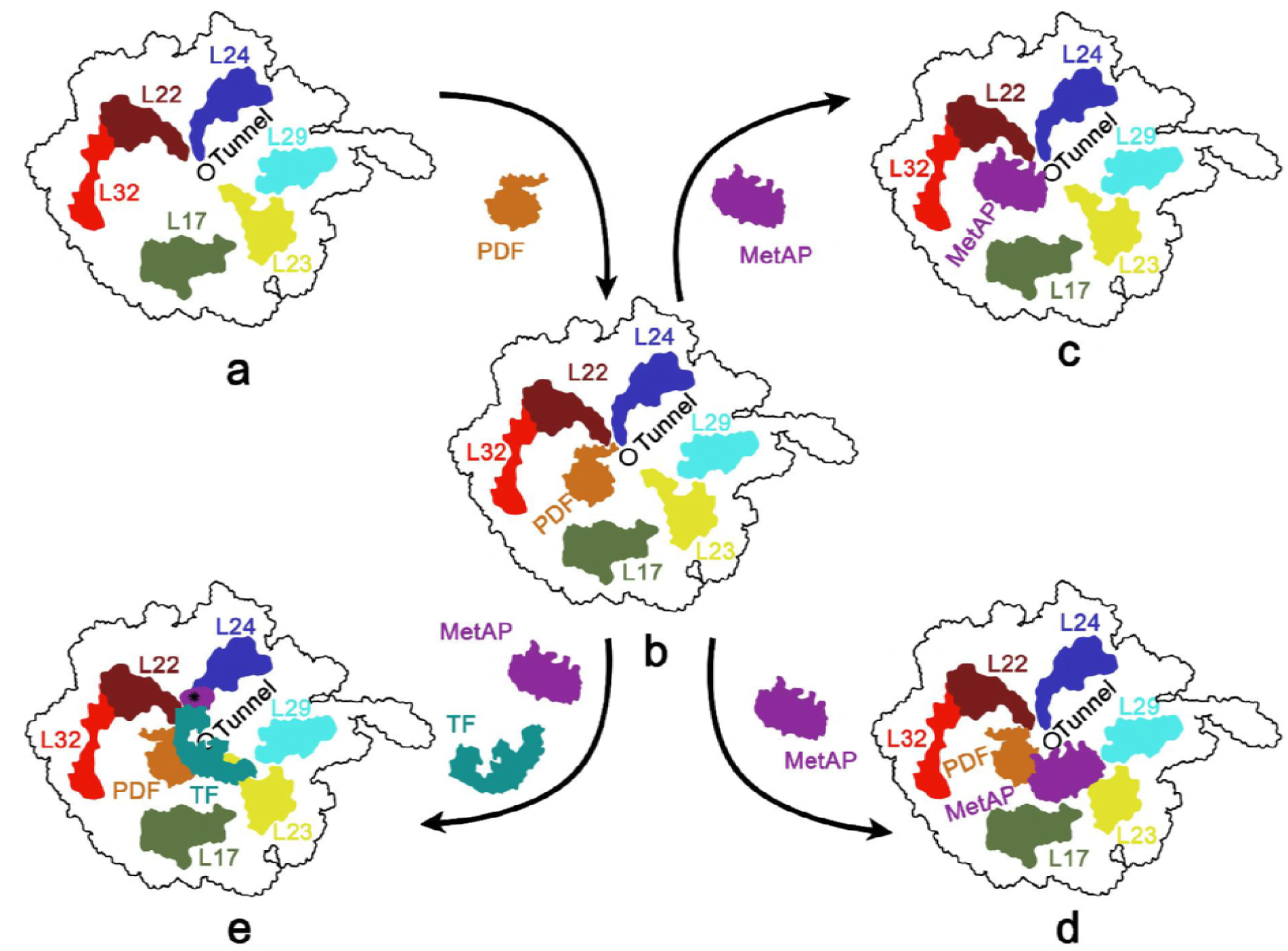
Schematic presentation of the proposed model for MetAP accessing ribosomal tunnel exit. During protein biogenesis, PDF is the first ligand to bind the 70S ribosome (a) near the tunnel exit at a specific site (b). Following the cleavage of formyl moiety from N-terminus of the emerging nascent polypeptide chain, it falls off the ribosome, emptying the surface for MetAP binding (c). While in simultaneous binding mode, MetAP anchors to an adjacent region of PDF near L23 where it primarily binds to rRNA and access the formyl free methionine to chop it off from the nascent chain (d). In presence of TF which binds to L23, MetAP relocates to yet another position near L24 (purple, marked with black star) and stays bound to the PPIase domain of TF.

## Materials and methods

### Purification of 70S ribosomes from E. coli

*E. coli* 70S ribosomes were purified following previously described protocol (Das et al, 1996) with minor modifications. MRE 600 cells from log phase were harvested and washed thrice with buffer A (20mM Tris-HCl pH 7.5, 10mM MgOAc_2_, 100mM NH_4_Cl, 5mM BME). Cells were resuspended in Buffer A and lysed using French press. The lysate was centrifuged (12,000g, 4°C, 20 min). The supernatant was collected and centrifugation was repeated until it stopped giving pellet. Crude ribosome was pelleted down (1,54,000g, 4°C, 2 hrs) from the supernatant and kept overnight in TMA 10 (Tris-HCl, pH 7.5, 10mM MgOAc_2_, 30mM NH_4_Cl,5mM BME). Pellet was resuspended in TMA 10 and homogenised for 1 hr in presence of 1M NH_4_Cl, after which the ribosomal preparation was centrifuged several times at 4°C at 15,000g till no pellet was seen. Supernatant solution was centrifuged at 4°C at 1,54,000g for 2 hr. The pellet was collected and kept overnight in TMA 10 and then resuspended. Sample was loaded on top of a 10%–40% linear sucrose gradient in TMA10 and centrifuged at 1,54,000g for 90 min at 4°C. Gradient was collected from bottom to top and relevant fractions were pooled together. The 70S ribosome fraction was concentrated in TMA 10 using 10KD Amicon Ultracentrifugal filter units (Millipore) and stored at -80°C.

### Expression and Purification of E. coli PDF

The gene encoding *E. coli* PDF was PCR amplified and subcloned into kanamycin resistant pET28a vector using primers 5’-CTGTTTTT<u>GCTAGC</u>ATGTCAGTTTTGCAAGTGTTACAT-3’ and 5’-GTGT<u>CTCGAG</u>TTAAGCCCGGGCTTTCAG-3’ containing NheI/XhoI restriction sites and transformed into BL21-DE3 strain. Transformed cells were grown in Luria-Bertan (LB) medium containing 0.5mg/mL kanamycin at 37°C until it reached an OD_600_ ~0.8. PDF expression was induced by 0.5mM IPTG, followed by an overnight incubation of cells at 27°C. Induced cells were harvested, washed and lysed in PDF buffer (50mM HEPES-KOH, pH 7.4, 100mM NaCl, 5mM NiCl_2_, 1mM TCEP, 2% glycerol). The cell lysate was centrifuged (18,000g, 45min, 4°C) and the overexpressed N-terminal His_6_-tagged PDF was purified with Ni Sepharose 6FF affinity column. The purified protein was stored at −80°C as glycerol stocks.

### Purification of E. coli MetAP

DNA encoding MetAP, type-1a, was amplified from *E. coli* genomic DNA and subcloned into pET28a vector encoding N-terminal poly-His-tag, using the primer set: 5′ primer: 5′-TATAA <u>G GATC C</u> ATGGCTATCTCAATCA- 3′; 3′ primer: 5′-TATAA <u>A AGCT T</u> TTATTCGTCGTGCGA- 3′ containing BamHI/HindIII restriction sites (underlined). *E. coli* BL21-DE3 cells were transformed with this clone and were grown to an OD_600_ of 1.2 at 37°C. Cells were induced with 1mM IPTG for 5 hours at the same temperature. Cells were harvested and dissolved in +TG buffer (50mM HEPES-NaOH, pH 8.0, 0.5M NaCl, 10% glycerol, 0.1% Triton-X 100, and 5mM imidazole) and lysed by sonication. The lysate was centrifuged at 18,000g for 20 min and the soluble fraction was filtered through 0.22μm pore size sterile filter. The filtered soluble fraction was loaded on top of the +TG pre-equilibrated Ni^+2^-NTA affinity flow column and the column was washed with the same buffer, followed by-TG buffer (50mM HEPES-NaOH, pH 8.0, 0.5M NaCl, and 5mM imidazole). The protein was eluted with elution buffer (50mM Hepes, pH 8.0, 0.5M NaCl, and 150mM imidazole). The protein was buffer exchanged in storage buffer (50mM HEPES, pH 8.0, 150mM NaCl) and concentrated using 10KD Amicon Ultra centrifugal filter units (millipore). All the buffer compositions were made following standard protocol (Addlagatta et al, 2005). The total yield of protein was approximately 266 mg for 2L of culture.

### Purification of E. coli Trigger Factor

BL21 cells were transformed with Pet28a vector containing gene encoding *E.coli* trigger factor protein. Transformed cells were grown in LB media at 37°C until it reached an OD_600_ ~0.8. Cells were induced with 0.4mM IPTG at 37°C for 4 hours. Cells were harvested, washed and dissolved in lysis buffer (50mM Phosphate buffer pH 8, 500mM NaCl, 1mM PMSF, 10mM Imidazole) and then lysed and centrifuged at 18000g (4°C) for 45min. The supernatant was collected and the His_6_ tagged TF were allowed to bind to Ni Sepharose resin for an hour. The protein bound resins were collected and washed repeatedly with lysis buffer. His_6_ tagged TF was then collected at 250mM Imidazole concentration. Purified protein was concentrated using 10KD Amicon Ultra centrifugal filter units (Millipore) in storage buffer (20mM Tris-HCl pH 7.5, 100mM NaCl, 1mM TCEP) and stored at −80°C as glycerol stocks.

### Gel electrophoresis and western blot analysis

Ribosome samples with or without bound proteins were loaded on a 12% denaturing SDS-PAGE gel and stained with Coomassie blue for visualizing protein bands. Recombinant proteins PDF, MetAP and TF were localized and estimated to have molecular weights of 20kDa, 30kDa and 55kDa respectively.

Western blot analysis was performed for ribosomes with or without the bound protein and for the recombinant protein peptide His_6_-tagged PDF (procured from GCC Biotech, India). Blots were probed with anti-His antibody. Binding of the secondary HRP-conjugated anti-rabbit antibodies (Millipore, USA) was analyzed using ImmunoCruz (Santa Cruz Biotechnology Inc., USA).

### Mass spectrometry analysis

Mass analysis was performed using 4800 MALDI TOF/TOF (model 4800, Applied Biosystems, USA) instrument operated in ‘reflectron’ mode. For MS/MS analysis, protein bands were excised from the gel and digested in gel using In-Gel-Tryptic-Digestion-Kit (Thermo Pierce). Mass analysis was performed using a saturated solution of CHCA (α-cyano-4-hydroxycinnamic acid) in 50% acetonitrile/0.1% trifluoroacetic acid. The MS/MS peaks of the most intense mass ions were searched against the NCBInr Database using MASCOT software (Matrix Science Ltd., London, UK). Peptides were matched to proteins when statistically significant MASCOT probability scores (<0.05) were consistent with experimental MW of the protein.

### Ribosome co-sedimentation assay

Ribosomes were incubated with a 60-fold excess of recombinant proteins for 30 mins at 37°C in binding buffers (20mM HEPES-KOH, pH 7.5, 100mM NH_4_OAc, 20mM MgCl_2_ and 1mM TCEP) and (20mM HEPES, pH7.5, 50mM KOAc, 15mM Mg(OAc)_2_, 0.1mM CoCl_2_) for PDF/TF and MetAP, respectively. Samples were pelleted through 5% sucrose cushion (45000g, 7 hrs, 4°C) and the supernatants and pellets were separately collected. Ribosomal pellets were re-suspended and the supernatants were dialysed and concentrated in binding buffers. Both were then resolved in a 12% SDS-PAGE gel. The absence of protein bands in the control lanes indicates that co-sedimentation of a protein occurs only due to the specific interaction of a protein with a ribosome. The binding of proteins was further validated by MSMS and was western blotted in case of PDF, since ribosomal proteins S4 and L4 were also of similar molecular weights as recombinant PDF.

### Sample preparation for cryo-EM imaging

Co-sedimentation (Sandikci et al, 2013) pellet of the 70S-PDF complex was dissolved in PDF binding buffer. 30-fold excess PDF was added and incubated with the complex (70S concentration 0.3μM) at 37°C for 30 minutes under constant shaking condition and then kept on ice until grid preparation.

60-fold excess purified MetAP was incubated with purified E. coli 70S ribosome (0.8 μM) in binding buffer (20mM HEPES, pH7.5, 50mM KOAc, 15mM Mg(OAc)_2_, 0.1mM CoCl_2_) at 37°C for 10 min and then incubated on ice.

To prepare the 70S-PDF-MetAP complex, 70S ribosome was incubated first with PDF (37°C, 30 mins) and MetAP was then added to the reaction mixture (37°C for 10 mins). The reaction mixture was kept on ice until grid preparation.

For preparation of the 70S-PDF-MetAP-TF complex, ribosome at concentration 0.55 μM was incubated with 60-fold excess PDF in binding buffer, followed by activated MetAP and then with TF. MetAP was activated by incubating in MetAP binding buffer at Co^2+^ concentration 0.05mM.

Cryo-EM grids of the 70S-protein complexes were prepared following standard procedures (Grassucci et al, 2007). Tedpella Lacey grids were used for 70S-PDF complexes with a blotting time of 6.5s. For all other complexes Tedpella Quantifoil grids were used with blotting time of 7s.

### Cryo-electron microscopy and 3D Image processing

Data were collected using 4Kx4K CCD camera with a physical pixel-size of 15 μm (corresponding to a pixel size of 1.89Å on the object scale) on a FEI (Eindhoven, The Netherlands) Tecnai G2 polara field emission gun electron microscope, equipped with low-dose kit and cryo-transfer holder at a total magnification of 78894X.

Data were primarily processed with SPIDER (Frank et al, 1996) package. Micrographs were evaluated after calculating power spectrum. Micrographs with defocus value in the range of 1 μm to 4 μm were checked for their Thon rings. Selected micrographs were grouped into defocus groups for CTF (Contrast Transfer Function) correction. Particles were selected based on a fast locally normalized correlation algorithm (Rath & Frank, 2004; Roseman, 2003). A background noise image was used to normalize the backgrounds of particle images.

Projections of a reference 3D map at uniform Euler angles were used to find different orientations of particle images. Auto-picked particles were low-pass filtered for better visualization during manual selection of good particles. Manually selected good particles were aligned to 2D projections of the 3D reference map, rotated at an interval of 15° Euler angle for each defocus groups. Aligned particles for each reference view were averaged to increase the signal to noise ratio. Particles were further selected based on the correlation cut-off threshold. The 3D reconstruction followed the standard SPIDER protocol with *E. coli* 70S ribosome filtered at very low frequency as a reference. Refinement of the 3D volume was done by iteratively adjusting the angles for finer resolution. CTF-correction was performed when the reconstructions from different defocus groups are merged. Supervised classification of the dataset was done as per the standard protocol (Gao et al, 2004). Fourier amplitudes of 3D volumes were enhanced using the x-ray scattering data. The 30S and 50S subunits and protein densities were isolated from the enhanced maps following the standard protocol of difference mapping. Reference-free 2D classification and unsupervised 3D classification by maximum likelihood method were performed on control (empty 70S ribosome) dataset to sort into ratcheted and non-ratcheted ribosomes (Schatz & van Heel, 1990; Scheres, 2012; van Heel, 1984; van Heel, 1990; van Heel & Frank, 1981). Pymol (Deleno Scientific) and Chimera (Pettersen et al, 2004) were used for cryo-EM map analysis and preparation of illustrations.

## Acknowledgements

This work was primarily supported by SERB, DST (India) sponsored project GAP327, and partially by CSIR Network project ‘UNSEEN’, CSIR-Indian Institute of Chemical Biology, Kolkata, India. We sincerely thank Prof. Marin van Heel for critical comments on the manuscript. We are grateful to Dr. Ilic Zoran (Wadsworth Center, NY, USA) for critically reviewing the manuscript. We acknowledge the supercomputing facility provided by CSIR Fourth Paradigm Institute (http://www.cmmacs.ernet.in). We thank Dr. Chandana Barat for providing TF clone, and Ms. Ravali and Mr. Saumya for assisting in preliminary cryo-EM data processing. Fund provided by NIPER for project students is gratefully acknowledged. SB and SA acknowledge CSIR, India, for senior research fellowship.

## Author contributions

JS, SB and SA designed and conceived experiments. SB and SA carried out bulk of the experiments, cryo-EM 3D image processing and analyzed the data. CB performed cryo-grid preparation, cryo-EM data collection and preliminary analysis of the datasets. JS wrote the paper with inputs from SA and SB.

